# Microfluidic delivery of cutting enzymes for fragmentation of surface-adsorbed DNA molecules

**DOI:** 10.1101/2021.03.31.437857

**Authors:** Julia Budassi, NaHyun Cho, Anthony Del Valle, Jonathan Sokolov

**Affiliations:** Department of Materials Science and Chemical Engineering, Stony Brook University, Stony Brook, New York, United States of America; Department of Physics and Astronomy, Stony Brook University, Stony Brook, New York, United States of America

## Abstract

We describe a method for fragmenting, in-situ, surface-adsorbed and immobilized DNAs on polymethylmethacrylate(PMMA)-coated silicon substrates using microfluidic delivery of the cutting enzyme DNase I. Soft lithography is used to produce polydimethylsiloxane (PDMS) gratings which form microfluidic channels for delivery of the enzyme. Bovine serum albumin (BSA) is used to reduce DNase I adsorption to the walls of the microchannels and enable diffusion of the cutting enzyme to a distance of 10mm. Due to the DNAs being immobilized, the fragment order is maintained on the surface. Possible methods of preserving the order for application to sequencing are discussed.

## Introduction

Significant progress in DNA sequencing has occurred over the last fifteen years, with dramatic improvement in throughput, in particular, as well as in haplotype phasing, read lengths and contig size [1-3]. Despite this, highly accurate and complete genome analysis at a reasonable cost and with rapid turnaround time such as would be desirable for personalized medicine has not yet been achieved. Short-read technologies (up to several hundreds of bases) are capable of generating Terabases of data but have difficulty in mapping structural variations and regions with long repeats. The ‘repeatome,’ comprising roughly half of the genome, has a role in gene expression and in disease and exhibits a relatively high rate of mutation [4]. Synthetic long-read techniques grafted onto the short-read platforms have provided improvement over the original methods [5-10] and some longer-read platforms have also appeared [11-16]. Nonetheless, no currently available technique is able to generate reads of a single DNA molecule greater than a few tens of kilobases. Since the range of human chromosome sizes is 47-249 Mbp, there is still a need to assemble relatively small sequenced fragments into contigs and any simplification of the process can have a significant impact.

All current sequencing requires the fragmentation of long DNA molecules into kilobase-sized pieces or smaller for analysis. Long-range positional order is lost for the currently-used methods. The most widely-used techniques are fragmentation by mechanical means or enzymatic mean [17]. The mechanical techniques include sonicaton, hydrodynamic shearing through orifices (driven by centrifugation or use of a syringe pump), focused acoustic shearing (commercialized by Covaris, Woburn, MA) and nebulization (DNA suspended in a shearing buffer which is forced through an orifice by compressed air or nitrogen gas). The enzymatic fragmentation methods are based on nicking enzymes, restriction enzymes or various transposons (such as Illumina’s Nextera system, which fragments and adds adapters in the same step, referred to as ‘tagmentation’). NEB has developed a product using a mixture of enzymes called ‘Fragmentase’ (New England Biolabs, Ipswich, MA). For all methods, to greater or lesser degree, there are issues of damage to the fragments and sequence bias of breaks in GC-rich vs. AT-rich regions [18].

It is clear that a method which preserves the sequential ordering of the fragments would be highly beneficial in simplifying the assembly problem. Two groups have published papers using localized cutting on surface-immobilized DNAs, one using atomic force microscopy to mechanically cut the molecules [19-21] while the second group used an electrochemical method to locally activate (with Mg^+2^ ions) enzymatic cutting [22]. This work, while highly interesting, involves cutting single (or very few) molecules at a time and is difficult to scale up. Our group has developed a method to use soft lithography stamps to allow cutting of significantly larger numbers of surface-immobilized DNAs in parallel [23]. In that work, DNAs are deposited onto a substrate by withdrawing a polymethylmethacrylate (PMMA)-coated silicon wafer out of a DNA solution, a technique that has been termed ‘molecular combing [24-26]. This method and a technique developed for optical mapping on surfaces [27], have been used to deposit DNAs of up to megabase pair length on flat substrates [28]. The DNAs are stretched, aligned and immobilized along the direction of sample withdrawal at densities that depend on solution concentration, buffer pH [29-30] and surface type. A soft lithography stamp [31], in the form of a polydimethylsiloxane (PDMS) grating produced from a silicon master (see Fig 1), is ‘inked’ with a DNase 1 solution and placed in contact with the surface containing the stretched and immobilized DNA molecules. The DNAs are cut at the contact points of the stamp, maintaining (on the surface) positional order. In that work [23], the DNAs were removed, en masse, by desorbing the DNA into buffer (NEBuffer 3.1, B7203S) at 75°C for 20 minutes or dissolving the substrate PMMA and purifying by phenol extraction. The fragments were end-repaired and sequenced using the PacBio platform (without amplification of the fragments in the case of desorption). Though the positional order was lost in those experiments, the cutting method was successfully demonstrated and some ideas for maintaining the order of the fragments were suggested.

**Fig 1.**
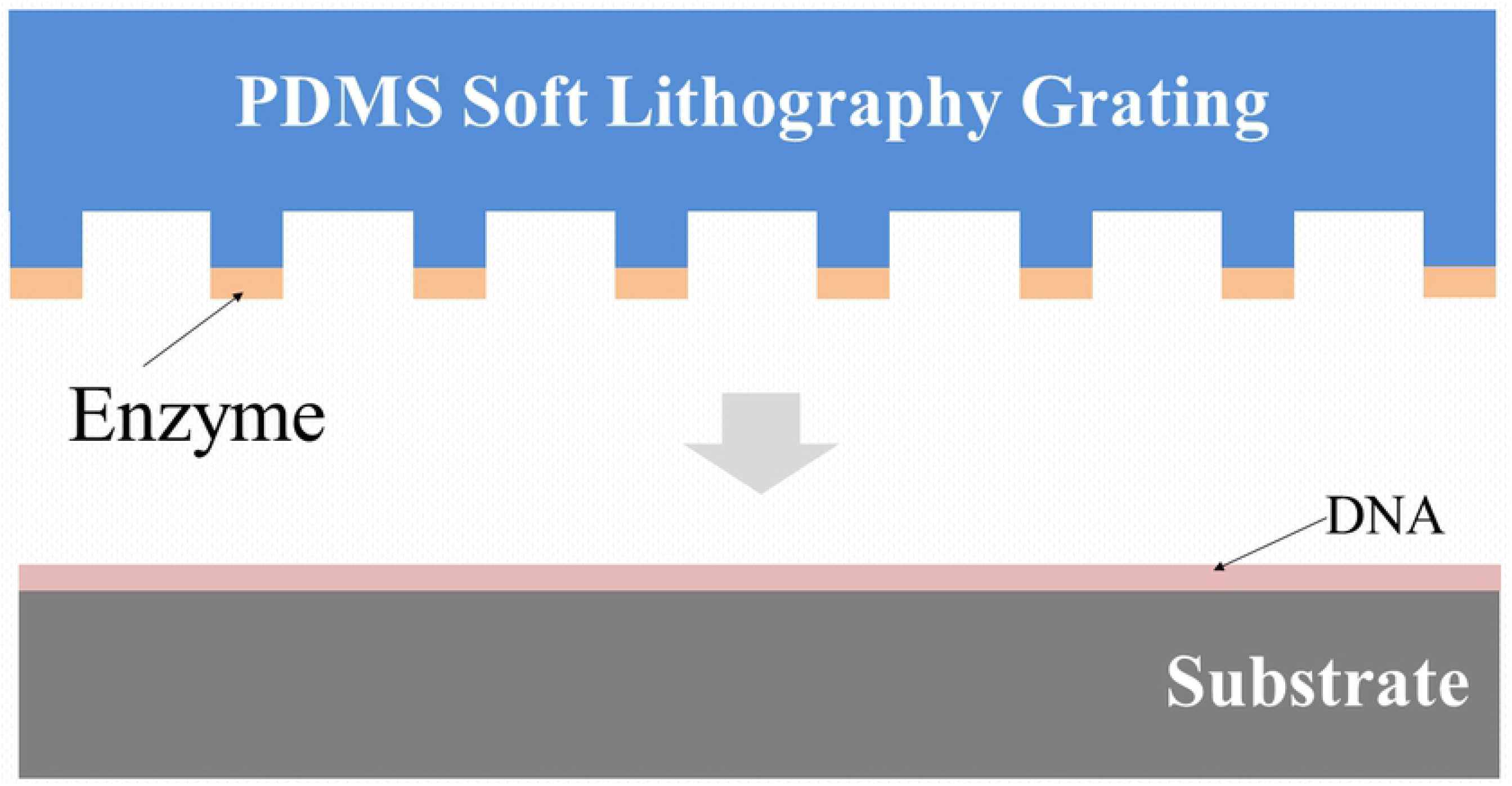
Schematic of stamping method for fragmenting surface-adsorbed. A PDMS stamp in the form of a grating is ‘inked’ with DNase I cuttting enzyme and is brought into contact with a surface on which DNA molecules have been deposited.

However, the inking method for delivering the DNase 1 enzymes is rather difficult to implement and we have sought to develop a more controllable technique. In this paper, we report on microfluidic delivery of the enzyme through micron-sized channels using soft lithography stamps. This technique is more reproducible and also lends itself to a variety of applications such as ordered removal of fragments or in-situ sequencing on the surface [32-33]. Another advantage of the method is that the application of the cutting enzyme is done from solution and so should have less steric hindrance than when applying by stamping.

## Materials and methods

### Sample preparation

Polished silicon wafers (Si(100), thickness 100-200μm thick, purchased from Wafer World, W. Palm Beach, FL) coated with PMMA layers, were used as substrates for DNA adsorption. The wafers were scribed and cleaved to make 1cm x 2cm samples. The wafers were cleaned using a modified Shiraki technique [34] as follows: (1) 10 minutes sonication in ethanol, (2) rinse in deionized (DI) water, (3) 15 minutes in boiling solution of 3:1:1 ratio (by volume) of water: ammonium hydroxide (28-30%): hydrogen peroxide (30%), (3) DI rinse, (4) 15 minutes in boiling solution of 3:1:1 ratio of water: sulfuric acid (98%): hydrogen peroxide (30%), (5) DI rinse, (6) one minute in 9:1 solution of water: hydrofluoric acid (49%), (7) DI rinse. The resulting surfaces were hydrophobic.

A 15 mg/ml solution of PMMA (molecular weight 70K, Polymer Source, Inc., Canada) in toluene was spun-cast (PWM32 spinner, Headway Research, Inc., Garland, Texas) onto the silicon wafers at 2500 RPM for 30 seconds. The thickness of the resulting films was measured using an ellipsometer (Auto El, Rudolph Research, Hackettstown, NJ) and was typically 70±8 nm. Following spin-coating, the samples were annealed for 1-4 hours at 130°C in an ion-pumped vacuum chamber (pressure ≤ 5 × 10^−7^ Torr) to remove adsorbed ambient and any remaining solvent.

DNA solutions for adsorption were produced in two steps. First, 200μl of a 50ng/μl solution (using Lambda DNA, New England Biolabs (NEB) N3011S), containing 1.5μl of the fluorescent dye SyBr Gold (Invitrogen, S11494, Thermo Fisher Scientific, Waltham, MA) was prepared in a buffer. The buffer was either a 6-12:50 mixture (by volume) of 0.1M sodium hydroxide: 0.02M 2-(n-morpholino) ethanesulfonic acid (MES) or 1X NEB DNase I reaction buffer (NEB B0303S, 1X is 10mM Tris-HCl, 2.5mM MgCl_2_, 0.5mM CaCl_2_). This solution was heated for one hour at 45°C to promote dye binding to the DNA. A further dilution in buffer by a factor of ten produced 2000μl of working solution at a DNA concentration of 5μl/mg.

DNA was adsorbed to the substrates by the technique called dynamic molecular combing [26]. The DNA solution is placed in a teflon well and the sample, held vertically with teflon tweezers, was lowered into the well and incubated for 30 seconds. The sample was then withdrawn at a rate of 1-2mm/s using a computer-controlled stepping motor attached to a linear drive stage (see Fig 2). The DNA molecules, preferentially attached by their ends, are stretched linearly and immobilized on the surface as they are removed from the solution (see Fig 2).

**Fig 2.**
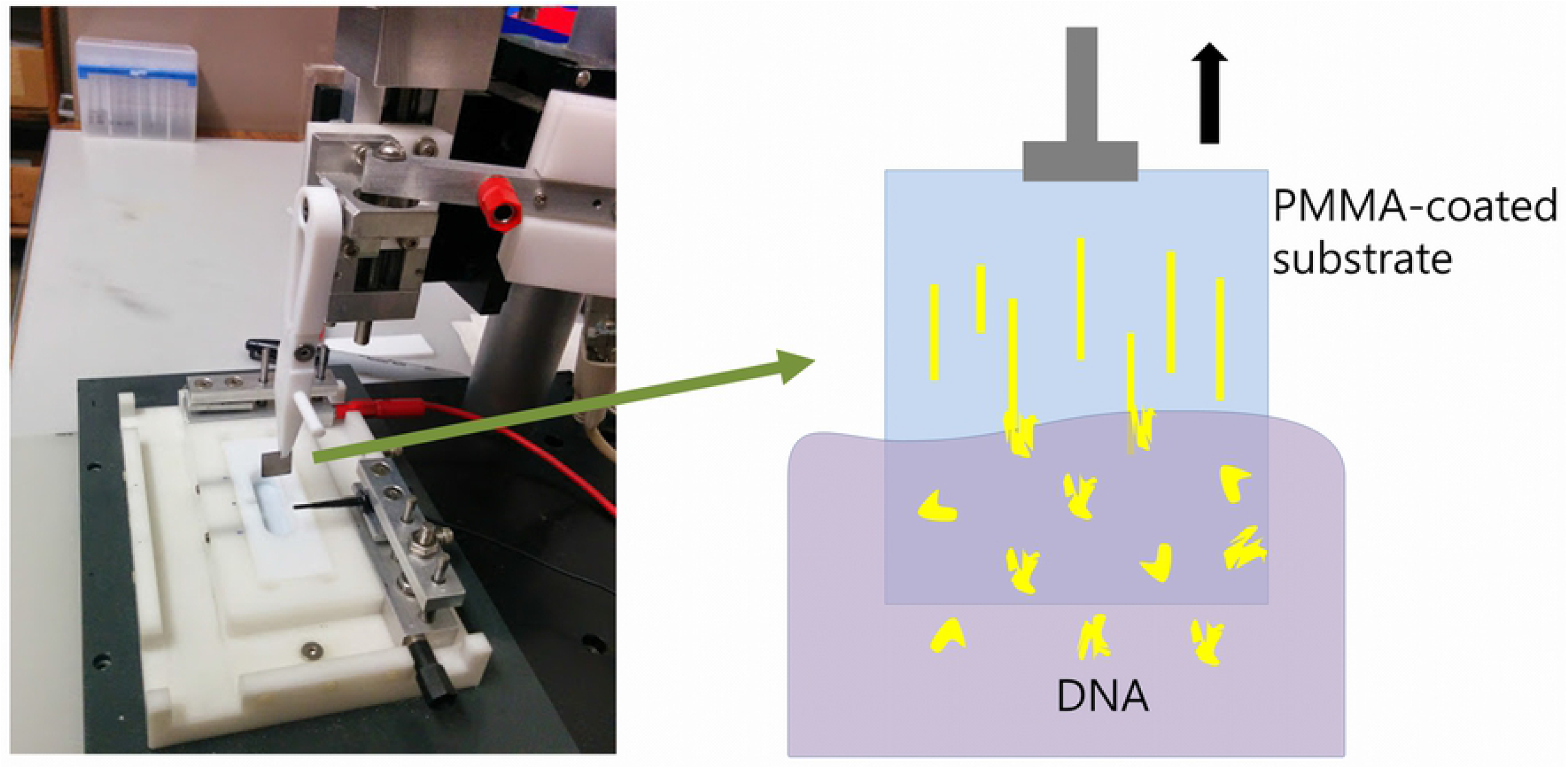
Apparatus for dip-coating (‘combing’) DNA molecules onto a substrate by withdrawal from solution.

### Production of PDMS microfluidic channels

The technique of soft lithography [31,35] was used to produce PDMS elastomer gratings. Silicon masters were made at the fabrication facilities of the Center for Functional Nanomaterials at Brookhaven National Laboratory. Firstly, a Cr/sodalime mask (aBeam Technologies,Hayward, CA) was used to used to pattern a photoresist-coated Si wafer of diameter 4” by UV exposure using a Karl Suss MA6 Mask Aligner (Suss MicroTec SE, Garching, Germany). The photoresist layer spun-cast onto the silicon wafers, nominally 1.1μm thick, was a positive resist, Shipley S1811 (Shipley Co., Marlborough, MA, USA). UV exposure was 5-40 seconds, followed by 110°C bake for 30s. The photoresist pattern was developed for 20-50s using a 2:3 mixture of MF-312 developer (Microposit, Rohm and Haas, Marlborough, MA) : water. Etching of the developed photoresist pattern to produce the silicon masters was done by reactive ion etching (RIE, Trion Phantom III RIEtcher, Trion Technology, Clearwater, FL, USA). The gas mixture was 40:10 SF_6_ : O_2_ at a pressure of 100mTorr. Etching power was 100-150 W and etching time was 300-700s. Leftover photoresist was dissolved in acetone. Optical microscopy (Olympus BH2 BHT) and atomic force microscopy (AFM, Digital Nanoscope 3000) were used to characterize the silicon patterns. Fig 3A shows an AFM image and Fig 3B the cross-section of a typical sample. The depth of the channels in the grating pattern was typically 2-5μm.

**Fig 3.**
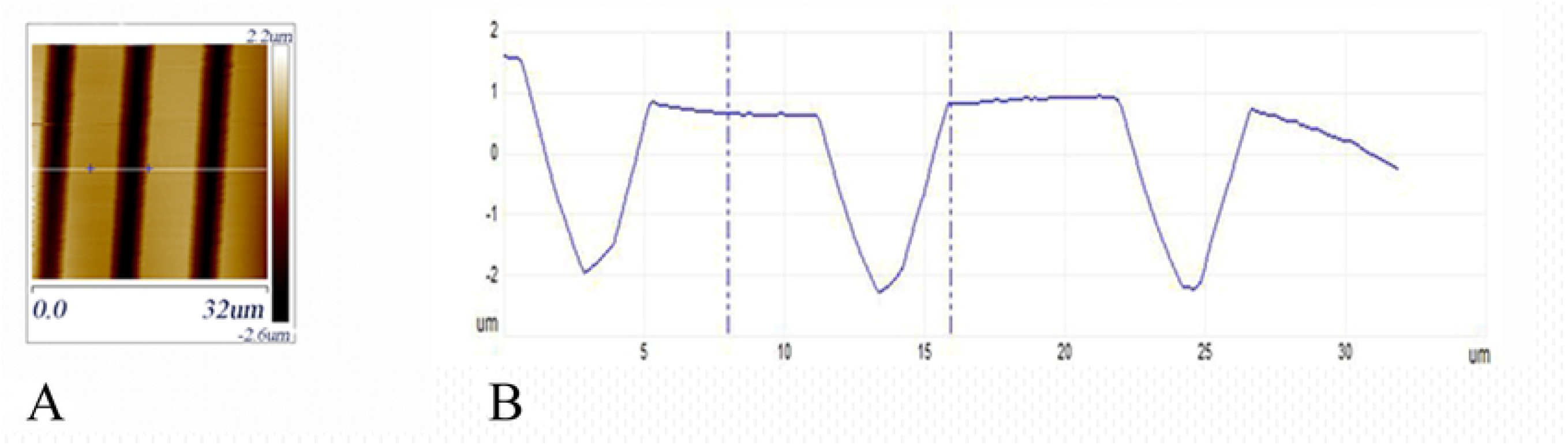
AFM image of silicon grating. (A) AFM topographical image of a silicon grating used as a master mold for making PDMS stamps. (B) Height cross-section along the white line in (A).

Soft lithography molds of PDMS were made using Sylgard 184 Silicone Elastomer (Dow Corning, Midland, MI, USA). A 10:1 mixture of elastomer and curing agent (by weight) was mixed thoroughly and trapped bubbles were removed by placing the mixture in a vacuum desiccator for one hour. The degassed PDMS was poured over the silicon mold to a thickness of approximately 5mm. The silicon mold was precoated with a thin film (less than 10nm) of PMMA, spun-cast from a 3mg/ml solution (molecular weight 70K). The purpose of the precoating was to reduce PDMS-silicon adhesion and facilitate removal of the PDMS layer. The PMMA-coated molds could be reused multiple times. The PMMA could also be removed with toluene and the wafers recoated for further use. The PDMS layers were cured at 60°C for 4 hours and then peeled off the molds. A typical cross-section of the grating, exposed by cutting the mold with a razor, is shown in the optical micrograph of Fig 4.

**Fig 4.**
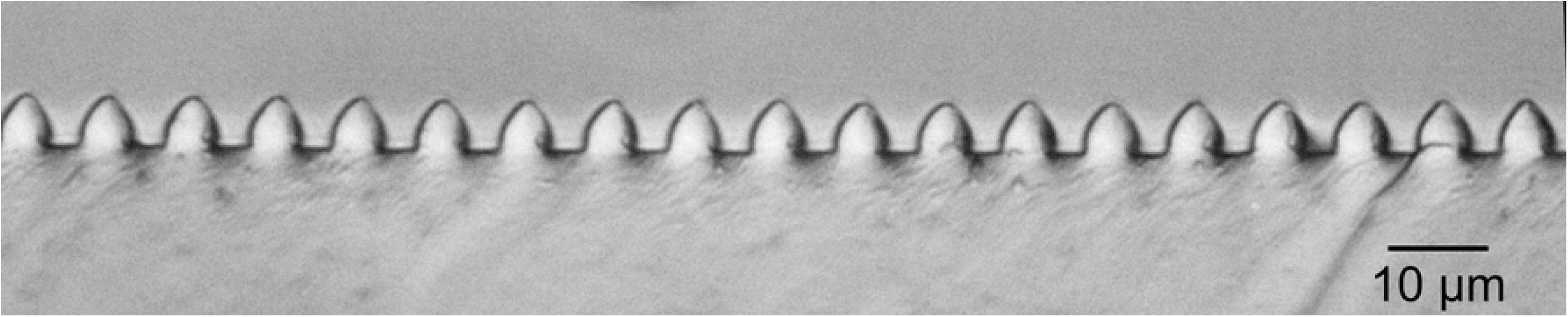
Optical Micrograph of a cross-section of a PDMS grating.

Microfluidic channels (approximately 4.5±0.1μm x 3.7±0.3μm x 12±2 mm, width by height by length, respectively) were made by placing the PDMS grating stamps in contact with the DNA-adsorbed substrates and tamping down the mold with tweezers to make good contact. An inlet/outlet hole of diameter 4mm had been previously cut through the PDMS layer using a biopsy punch (Integra, Miltex, Princeton, NJ USA)) and a liquid reservoir (also made from PDMS) with inner diameter of 6mm and height of 25mm was sealed to the stamp above the hole with PDMS (painted on and cured) (see Fig 5). The far end of the channels, away from the inlet/outlet, was sealed with PDMS, producing closed end channels. The cutting enzyme, here DNase I (NEB B0303S, Ipswich, MA USA), is delivered through the channels, as described below. The DNase I cuts the surface-immobilized DNAs along the channels while the PDMS stamp protects the DNA between the channels from being cut.

**Fig 5.**
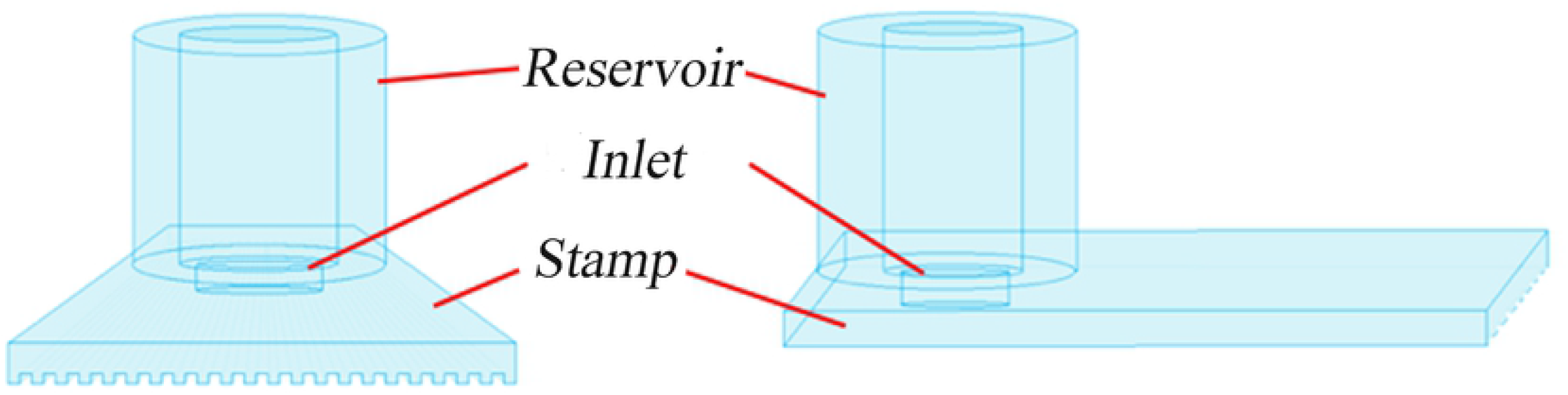
End-on and side views of a PDMS grating appended fluid reservoir.

### Results and discussion

Preliminary to doing the patterned cutting of DNA, we prepared solutions at different concentrations of DNase I and placed 3μl drops onto PMMA-coated samples with adsorbed DNA. The samples were heated at 40°C for 20 minutes, with the drops covered by mineral oil (M5904, MilliporeSigma, Burlington, MA) to prevent evaporation. They were then imaged by fluorescence microscopy to determine an effective enzyme concentration for cutting. The stock DNase I solution of 2Units(U)/μl was diluted in DNase I Reaction Buffer to concentrations 0.024U/μl, 0.048U/μl and 0.095U/μl (the recommended concentration for reactions in solution is given by the manufacturer as 0.02U/μl). Effective digestion was found for both of the higher concentrations, though somewhat more completely for the highest concentration. (see Fig 6). In further experiments, the concentration of 0.095U/μl was used unless noted otherwise. These results are consistent with the work of Gueroui et al [36], who observed digestion of combed DNA on a PMMA surface under similar conditions. (They also observed that for the restriction endonucleases HindIII and DraI the solution-level biochemical activity was not observed. We found the same result for PvuI.)

**Fig 6.**
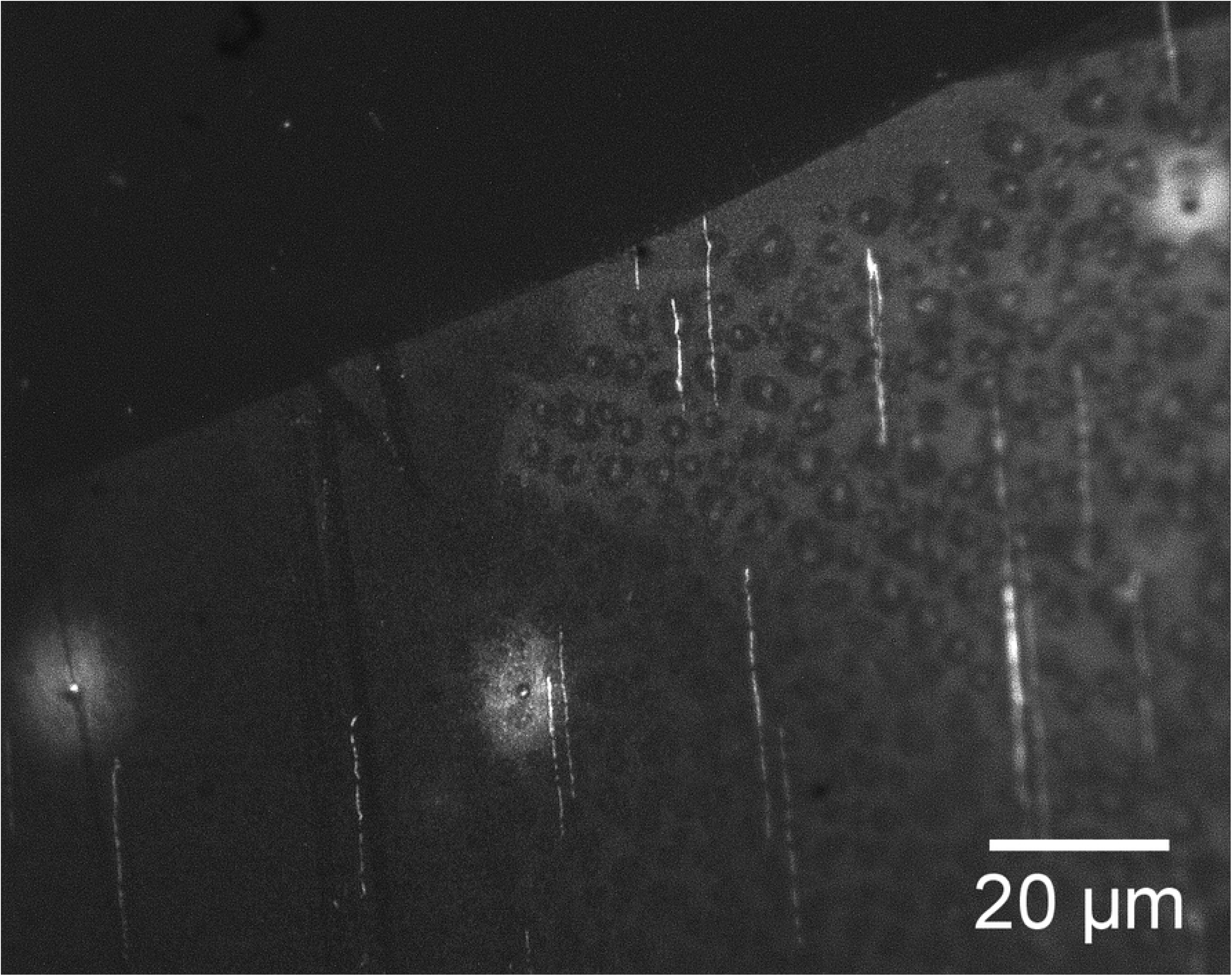
Fluorescence image of SyBr Gold labeled DNA. Upper left area was covered with a solution containing 0.095U/μl of DNase I in NEB DNase I Reaction Buffer and shows effective digestion of DNA in that region.

For the first set of cutting experiments, a PDMS stamp placed in contact with a DNA sample had its reservoir filled up with 300μl of the DNase I solution. To fill the long, narrow microfluidic channels (micron-sized cross-section by mm lengths) with the solution can be done in a number of ways—using capillary action (the PDMS surface needs to be made hydrophilic), applying vacuum at an open end away from the reservoir or applying pressure above the liquid in the reservoir, for example. We have used a convenient method [37], termed by the authors the ‘channel outgas technique.’ In this method, pressure is lowered above the reservoir (or the entire device is submerged in the filling liquid), causing air bubbles from the channels to escape through the liquid due to the buoyancy effect and allowing the channels to be filled with solution from the reservoir. The sample with stamp and reservoir was placed in a vacuum chamber (using an Edwards diaphragm pump having a teflon-coated diaphragm to enable pumping of high vapor pressure liquids) and the pressure was lowered to 20 Torr for 40 minutes. The sample, with channels now filled with the enzyme solution, was removed from the vacuum chamber and placed on a 40°C hotplate for 90 minutes to effect DNA digestion in the channels. The result was that digestion only occurred close to inlet of the reservoir, to a distance of less than 0.1mm. This raised a concern that perhaps the DNase I enzyme was damaged due to shearing forces exerted during the filling [38]. Therefore, it was decided to fill the channels first with buffer as above (20 Torr for 40 minutes) and then to add enzyme solution to the reservoir and allow penetration into the channels by diffusion (at 40°C for 90 minutes) through the liquid. The resulted in effective cutting of the DNA to a distance of 1.1±0.2mm from the inlet (Fig 7).

**Fig 7.**
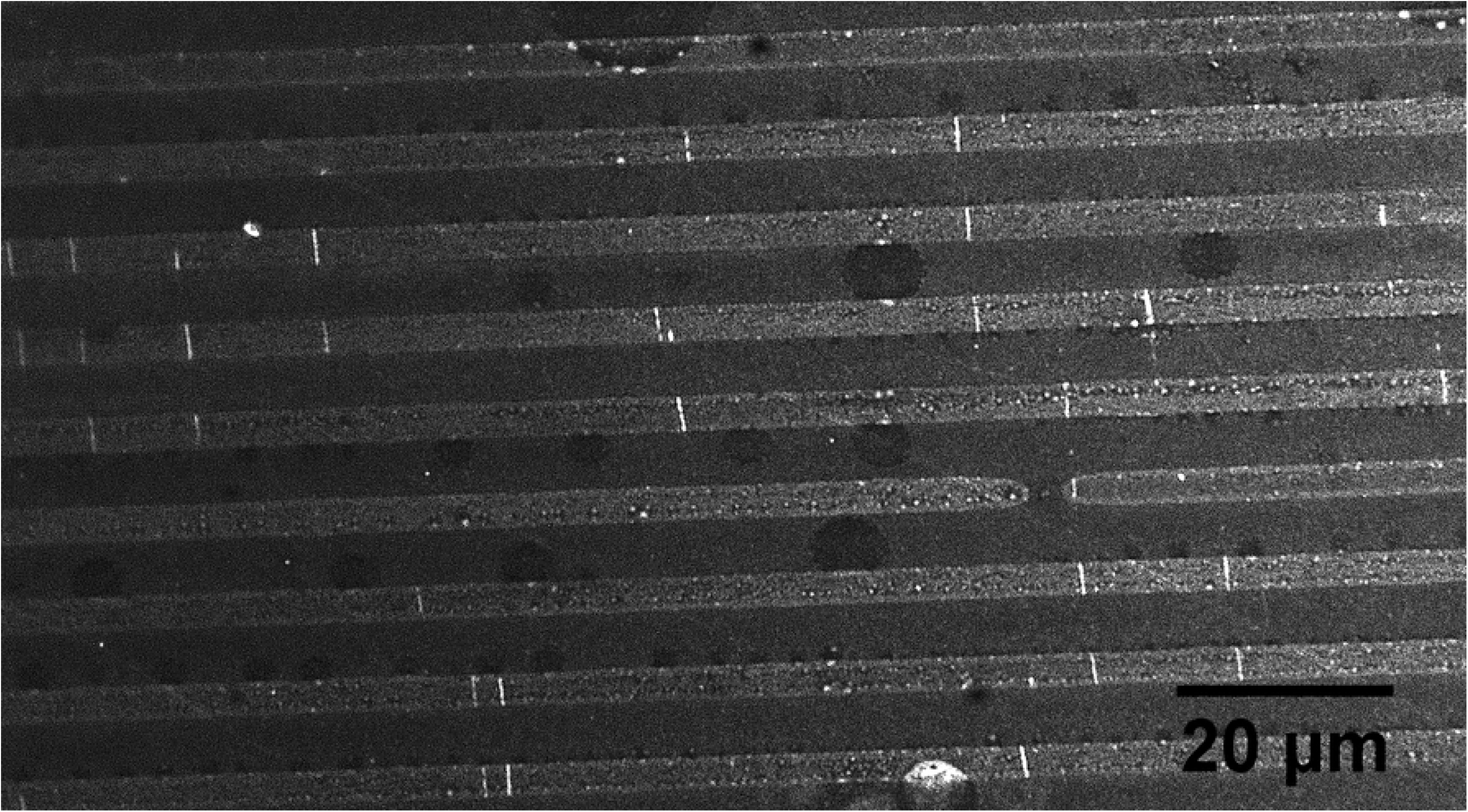
Fluorescence image of fragmented DNA remaining after digestion by DNase I diffusing through microfluidic channels. Distance from reservoir inlet is 1.1mm.

Following this modest improvement, we tried a series of similar experiments, lengthening the time of the heating/diffusion step up to 5 hours at 40°C. Experiments for times of 1.5, 2, 3, 4 and 5 hours all showed cutting up to a distance of approximately 1mm. A set of experiments varying the concentration of DNase I was tried next, with concentrations of 0.195, 0.295, 0.395, 0.495 and 0.595 U/μl used (vacuum fill of buffer as above, followed by 2 hour 40°C heating/diffusion step). No clear trend was discernible, though the best sample, for 0.495U/μl, had a cutting distance of 1.8mm.

At this point, it occurred to us that enzyme adsorption to the walls of the channels might be limiting the diffusion of the DNase I. Previous studies [39,40] have shown that proteins may be adsorbed to PDMS and also that bovine serum albumin (BSA) may be used to block protein adsorption [41]. Two experiments were conducted in which the vacuum filling of the channels with buffer was followed by a heating/diffusion step of 1 hour at 40°C with the reservoir filled with a solution containing both DNase I (at 0.096U/μl) and BSA (NEB B9000S) at 0.13mg/ml or 0.40mg/ml. The lower BSA concentration had little effect on the cutting distance. However, the higher concentration sample showed enzymatic cutting to a distance of 3.3mm.

Next, we decided to try to diffuse in the BSA separately from, and before, the cutting enzyme. In addition, due to the sometimes excessive bubbling of the liquid in the reservoir during vacuum filling (the boiling point of water at 20 Torr is 21.9°C, quite close to typical room temperature), the vacuum filling was done at 120 Torr for 40 minutes. The sample was also tilted at 45° to the horizontal to promote escape of gas bubbles from the channels. Following the vacuum filling, BSA was added to the reservoir to a concentration of 0.40mg/ml and left to incubate at 40°C for 1 hour. (As a check on the diffusion rates of BSA, we ran tests using fluorescently-labeled FITC-BSA (ThermoFisher Scientific, Waltham, MA), see Fig 8.) Afterwards, DNase I was added to 0.095U/μl in the reservoir and incubated for 2 hours. With these changes, the effective cutting distance was increased to 5.0±0.4mm.

**Fig 8.**
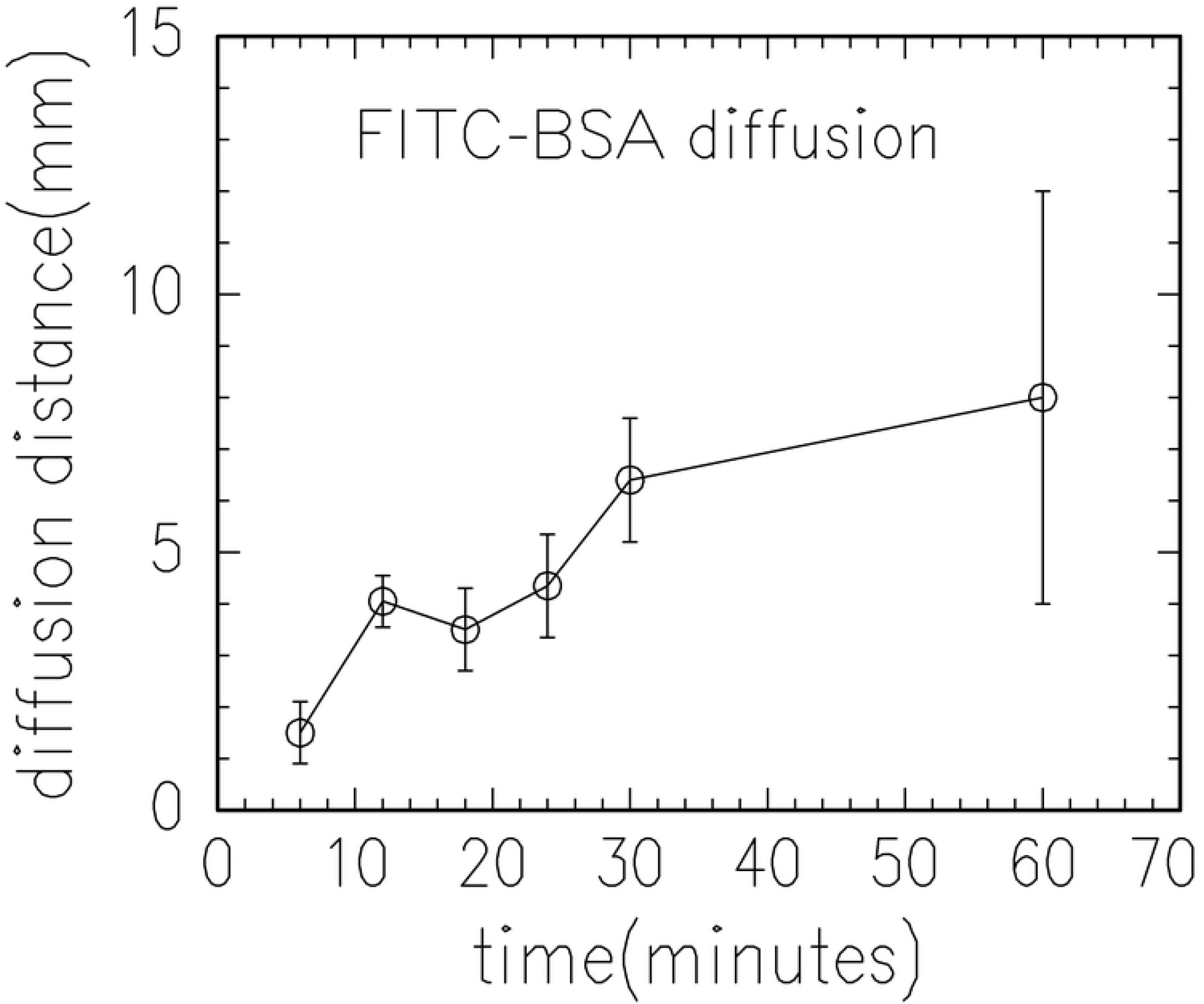
Measured diffusion distance of FITC-labeled BSA through micorfluidic channels versus diffusion time.

Further optimization was obtained by using the same steps but varying the BSA concentration up to 1.07 mg/ml, varying BSA incubation time up to 3 hours and DNase I incubation time also up to 3 hours. This produced a significant improvement, with cutting distances of 10mm consistently achieved. Fig 9 shows a sample with effective digestion for a sample with high density of adsorbed DNA.

**Fig 9.**
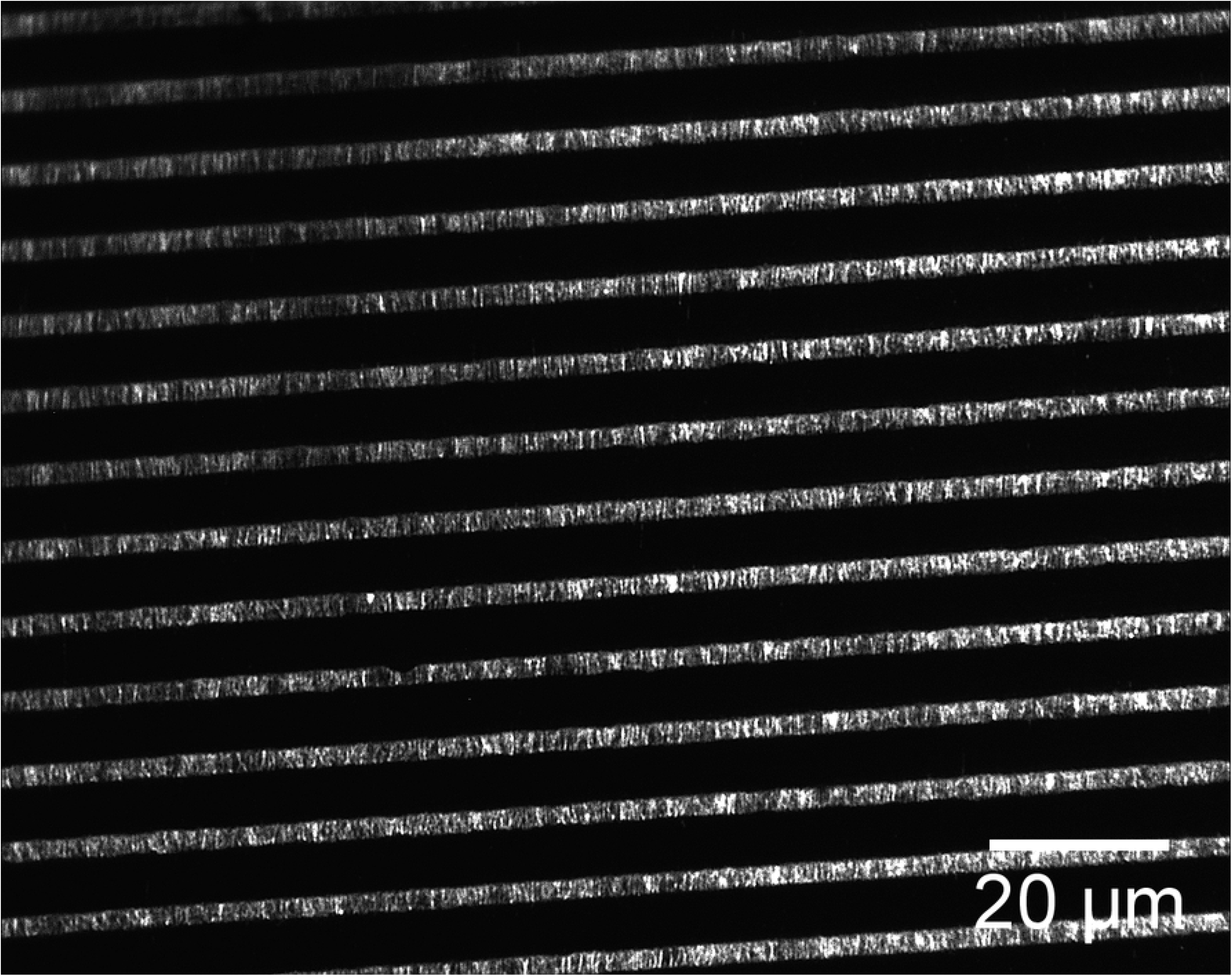
DNA (at high density) fragmented on a surface by DNase I. Distance from inlet is 8.7mm.

Finally, an effort was made to shorten the times of the various steps, keeping the optimum concentrations fixed. The following streamlined protocol was found:

1. 15 minutes of vacuum filling of the channels at a pressure of 120 Torr. The reservoir was filled with 150μl of DNase I Reaction Buffer.
2. The sample was placed on a 40°C hotplate. 8μl of BSA stock was added to the reservoir, making the BSA concentration 1.07mg/ml. The solution was mixed by careful up-and-down pipetting.
3. 60 minutes incubation with reservoir covered by parafilm to reduce evaporation.
4. Addition of 7.5μl of DNase I stock to reservoir, making the concentration 0.09Units/μl. Follow with mixing by careful up-and-down pipetting.
5. 1 hour of incubation at 40°C with reservoir covered by parafilm.

## Conclusions

We have demonstrated an effective and reproducible method for the fragmentation of surface-adsorbed and immobilized DNAs using soft lithography and microfluidic delivery of an anti-fouling coating (BSA) and the cutting enzyme (DNase I). This method also lends itself to ordered microfluidic removal of the fragments for sequencing applications or in-situ Next Generation Sequencing [33]. Removal of the fragments is complicated by the competing requirements of having relatively strong DNA-surface interactions, to enable immobilization on the surface, versus needing relatively weak interactions to allow desorption. One approach, which we are currently exploring, is to use a substrate which exhibits a reversible solubility switch from water-soluble to water-insoluble [42]. The immobilization is done in the water (and DNA-compatible buffer)-insoluble state while desorption is done into a water-based buffer which may be flowed through the channels. Alternatively, rather than use long microchannels, a PDMS stamp with holes used to create separate chambers could be used for fragmenting and amplifying/sequencing in-situ. Also, as noted above, steric hindrances can affect enzyme activity and the use of microporous substrates [43,44] can broaden the range of useable enzymes.

## Acknowledgments

This research used resources of the Center for Functional Nanomaterials, which is a U.S. DOE Office of Science Facility, at Brookhaven National Laboratory under Contract No. DE-SC0012704. We thank Ke Zhu, Donald Liu, Alan Gan, Sara Goodwin and Adriana Pinkas-Sarafova for assistance with the experiments and for discussions.

## References

1. Mardis, ER. Next-generation DNA sequencing methods. Annual Review of Genomics and Human Genetics. 2008; 9: 387–402.

2. Goodwin, S, McPherson, JD, McCombie, WR. Coming of age: ten years of Next-Generation Sequencing Technologies. Nature Reviews Genetics. 2016; 17: 333–351.

3. Giani, AM, Gallo, GR, Gianfranceschi, L, Formenti, G. Long walk to genomics: History and current approaches to genome sequencing and assembly. Computational and Structural Biotechnology Journal. 2020; 18: 9–19.

4. Hannan, AJ. Tandem repeats mediating genetic plasticity in health and disease. Nature Review Genetics. 2018; 19(5): 286–298.

5. Kitzman, JO, Mackenzie, AP, Adey, A, Hiatt, JB, Patwardhan, RP, Sudmant, PH, et al. Haplotype-resolved genome sequencing of a Gujarati Indian individual. Nature Biotechnology. 2011; 29: 59–63.

6. Peters, BA, Kermani, BG, Sparks, AB, Alferov, O, Hong, P, Alexeev, A, et al. Accurate whole-genome sequencing and haplotyping from 10 to 20 human cells. Nature. 2012; 487: 190–195.

7. Amini, S, Pushkarev, D, Christiansen, L, Kostem, E, Royce, T, et al. Haplotype-resolved whole-genome sequencing by contiguity-preserving transposition and combinatorial indexing. indexing indexing. Nature Genetics. 2014; 46: 1343–1349.

8. Zheng, GX, Lau, BT, Schnall-Levin, M, Jarosz, M, Bell, JM, et al. Haplotyping germline and cancer genomes with high-throughput linked-read sequencing. Nature Biotechnology. 2016; 34: 303–311.

9. Bruinsma, S, Burgess, J, Schlingman, D, et al. Bead-linked transposomes enable a normalization-free workflow for NGS library preparation. BMC Genomics. 2018; 19: Article 722.

10. Chen, H, Yao, J, Fu, Y, Pang, Y, Wang, J, Huang, Y. Tagmentation on microbeads: restore long-range DNA sequence information using Next Generation Sequencing with library prepared by surface-immobilized transposomes. ACS Applied Materials and Interfaces. 2018; 10: 11539–11545.

11. Eid, J, Fehr, A, Gray, J, Luong, K, Lyle, J, et al. Real-time DNA sequencing from single polymerase molecules. Science. 2009; 323(5910): 133–138.

12. Deamer, D, Akeson, M, Branton, D. Three decades of nanopore sequencing. Nature Biotechnology. 2016; 34: 518–524.

13. Gordon, D, Huddleston, J, Chaisson, MJ, Hill, CM, Kronenberg, ZN, Munson, KM, Malig, M, Raja, A, Fiddes, I, Hillier, LW, Dunn, C, Baker, C, Armstrong, J, Diekhans, M, Paten, B, Shendure, J, Wilson, RK, Haussler, D, Chin, CS, Eichler, EE. Long-read sequence assembly of the gorilla genome. Science. 2016; 352(6281): aae0344-1 to -7.

14. Mantere, T, Kersten, S, Hoischen, A. Long-read sequencing emerging in medical genetics. Frontiers in Genetics. 2019; 10: Article 426.

15. Shafin, K, Pesout, T, Lorig-Roach, R, et al. Nanopore sequencing and the Shasta toolkit enable efficient de novo assembly of eleven human genomes. Nature Biotechnology. 2020; https://doi.org/10.1038/s41587-020-0503-6.

16. Amarasinghe, SL, Su, S, Dong, X, Zappia, L, Ritchie, ME, Gouil, Q. Opportunities and challenges in long-read sequencing data analysis. Genome Biology. 2020; 21(1): 1–16.

17. Knierim, E, Luckel, B, Schwarzl, J. M, Schuelke, M, Seelow, D. Systematic comparison of three methods for fragmentation of long-range PCR products for Next Generation Sequencingl. PloS ONE. 2011; 6(11): e28240.

18. Poptsova, M, Il’icheva, I, Nechipurenko, D, et al. Non-random DNA fragmentation in next-generation sequencing. Science Reports. 2015; 4: 4532 https://doi.org/10.1038/srep04532.

19. Kurosawa, O, Okabe, K, Washizu, M. DNA analysis based on physical manipulation Proceedings of the 13th Annual International Conference on Micro Electro Mechanical Systems (MEMS 2000) pp. 311–316.

20. Yamamoto, T, Kurosawa, O, Kabata, H, Shimamoto, N, Washizu, M. Molecular surgery of DNA based on electrostatic micromanipulation. IEEE Transactions on Industry Applications. 2000; 36(4): 1010–107.

21. Kurosawa, O, Washizu, M. Dissection, acquisition and amplification of targeted position of electrostatically stretched DNA. Journal of Electrostatics. 2007; 65: 423–430.

22. Mizuno, A, Katsura, S. DNA immobilized on glass, localized Mg ions activated EcoRI restriction enzyme manipulation of a large DNA molecule using the phase transition. Journal of Biological Physics. 2002; 28: 587–603.

23. Cho, N, Goodwin, S, Budassi, J, Zhu, K, McCombie, WR, Sokolov, J. Fragmentation of surface adsorbed and aligned DNA molecules using soft lithography for Next-Generation Sequencing. Journal of Biosensors & Bioelectronics. 2017; 8(3): DOI: 10.4172/2155-6210.1000247.

24. Bensimon, A, Simon, A, Chiffaudel, A, Croquette, V, Heslot, F, Bensimon, D. Alignment and sensitive detection of DNA by a moving interface. Science. 1994; 265(5181): 2096–8.

25. Bensimon, D, Simon, AJ, Croquette, V, Bensimon, A. Stretching DNA with a receding meniscus: experiments and models. Physical Review Letters. 1995; 74(23): 4754–4757.

26. Michalet, X, Ekong, R, Fougerousse, F, Rousseaux, S, Schurra, C, Hornigold, N, van Slegtenhorst, M, Wolfe, J, Povey, S, Beckmann, JS, Bensimon, A. Dynamic molecular combing: stretching the whole human genome for high-resolution studies. Science. 1997; 277(5331): 1518–1523.

27. Schwartz, DC, Li, X, Hernandez, LI, Ramnarain, SP, Huff, EJ, Wang, Y-K. Ordered restriction maps of Saccharomyces cerevisiae chromosomes constructed by optical mapping. Science. 1993; 262(5130); 110–114.

28. Kaykov A, Taillefumier T, Bensimon A, Nurse P. Molecular combing of single DNA molecules on the 10 megabase scale. Scientific Reports. 2016; 6:19636: DOI:10.1038/srep19636.

29. Allemand, JF, Bensimon, D, Jullien, L, Bensimon, A, Croquette, V. pH-dependent specific binding and combing of DNA. Biophysical Journal. 1997; 73: 2064–2070.

30. Benke, A, Mertig, M, Pompe, W. Ph- and salt-dependent molecular combing of DNA: experiments and phenomelogical model. Nanotechnology. 2010; 22(3): doi: 10.1088/0957-4484/22/3/035304.

31. Xia, Y, Whitesides, GM. Soft lithography. Angewandte Chemie International Edition. 1998; 37: 550–575.

32. Schwartz, J, Lee, C, Hiatt, JB, et al. Capturing native long-range contiguity by in-situ library construction and optical sequencing. Proceedings of the National Academy of Science. 2012; 109(46): 18749–18754.

33. Feng, K, Costa, J, Edwards, JS. Next-generation sequencing library construction on a surface. BMC Genomics. 2018; 19:416: https://doi/10.1186/s12864-018-4794-4.

34. Okumura, H, Akane, T, Tsubo, Y, Matsumoto, S. Comparison of conventional surface cleaning methods for Si molecular beam epitaxy. Journal of the Electrochemical Society. 1997; 144: 3765–3768.

35. Whitesides, GM, Ostuni, E, Takayama, S, Jiang, X, Ingber, DE. Soft lithography in biology and biochemistry. Annual Review of Biomedical Engineering. 2001; 3: 335–373.

36. Gueroui, Z, Place, C, Freyssingeas, E, Berge, B. Observation by fluorescence microscopy of transcription on single combed DNA. Proceedings of the National Academy of Sciences. 2002; 99(9): 6005–6010.

37. Monahan, J, Gewirth, AA, Nuzzo, RA. Method for filling complex polymeric microfluidic devices and arrays. Analytical Chemistry. 2001; 73(13): 3193–3197.

38. Clarkson, JR, Cui, ZF, Darton, RC. Protein denaturation in foam. Journal of Colloid and Interface Science. 1999; 215(2): 333–338.

39. James, CD, Davis, RC. Kam, L, Craighead, HG, Isaacson, M, Turner, JN, Shain, W. Patterned protein layers on solid substrates by thin stamp microcontact printing. Langmuir. 1998; 14(4): 741–744.

40. Bernard, A, Delamarche, E, Schmid, H, Michel, B, Bosshard, HR, Biebuyck, H. Printing patterns of proteins. Langmuir. 1998; 14(9): 2225–2229.

41. Ostuni, E, Chen, CS, Ingber, DE, Whitesides, GM. Selective deposition of proteins and cells in arrays of microwells. Langmuir. 2001; 17: 2828–2834.

42. Linder, V, Gates, BD, Ryan, D, Parviz, BA, Whitesides, GM. Water-soluble sacrificial layers for surface micromachining. Small. 2005; 1(7): 730–736.

43. Fuke A, Suzuki, T, Nakama, K, Kabata, H, Kotera, H. High-throughput gene analysis using suspending DNA fibers (SDFs) on a micro glass-phonorecord. Twelfth International Conference on Miniaturized Systems for Chemistry and Life Sciences. October 12 - 16, 2008; San Diego, California, USA, 1519–1521.

44. Trigo-Lopez, M, Vallejos, S, Reglero Ruiz, JA, Ramos, C, Beltran, S, García, FC, García, JM. Fabrication of microporous PMMA using ionic liquids: An improved route to classical ScCO2 foaming process. Polymer. 2019; 183: 121867.

